## Supplemental data including 12 figures for "Metabolic reprogramming of cancer cells by JMJD6-mediated pre-mRNA splicing is associated with therapeutic response to splicing inhibitor"

Figure S1-12

Table S1-8

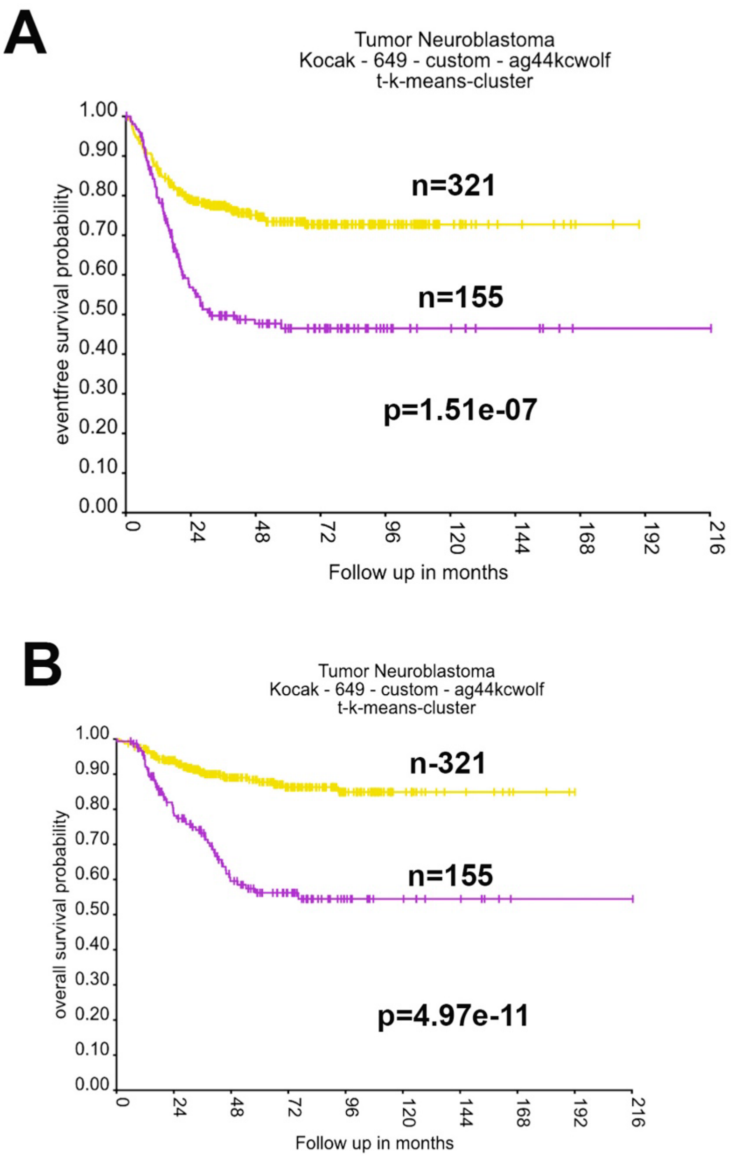


**Figure S1. Association of 17q essential genes with patient survival.** (**A**) Kaplan-Meier survival curve showing 17q essential gene signature is correlated with a worse event-free survival (Kocak dataset, GSE45547). (**B**) Kaplan-Meier survival curve showing 17q essential gene signature is correlated with a worse overall survival (Kocak dataset, GSE45547).

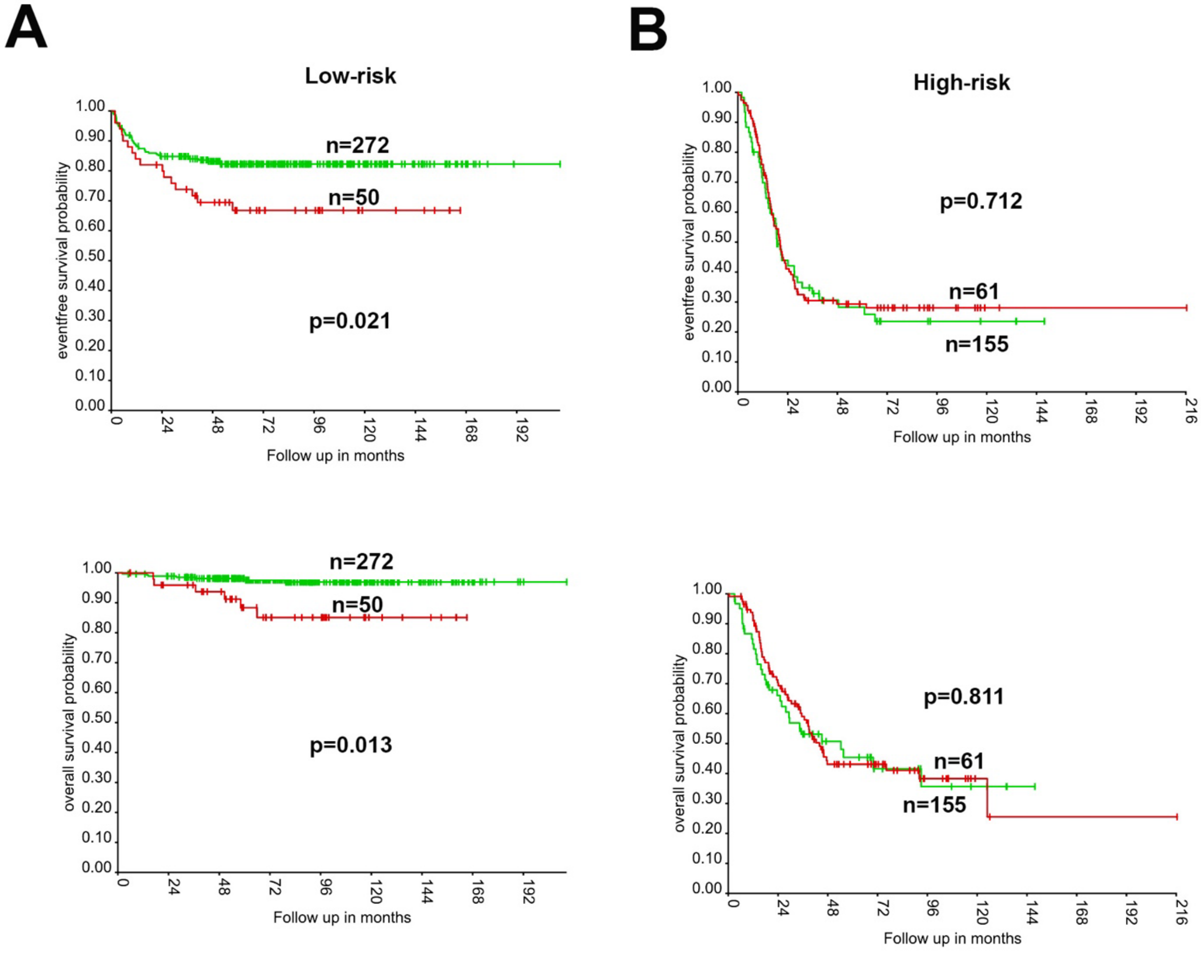


**Figure S2. Association of 17q essential genes with survival in low-risk and high-risk patients. (A)** Kaplan-Meier survival curve showing that the expression of 17q essential gene signature is correlated with worse event-free survival (top) and overall survival (bottom) in the low-risk patients (SEQC dataset). **(B)** Kaplan-Meier survival curve showing that the expression of 17q essential gene signature has no correlation with event-free survival (top) and overall survival (bottom) in the high-risk patients (SEQC dataset).

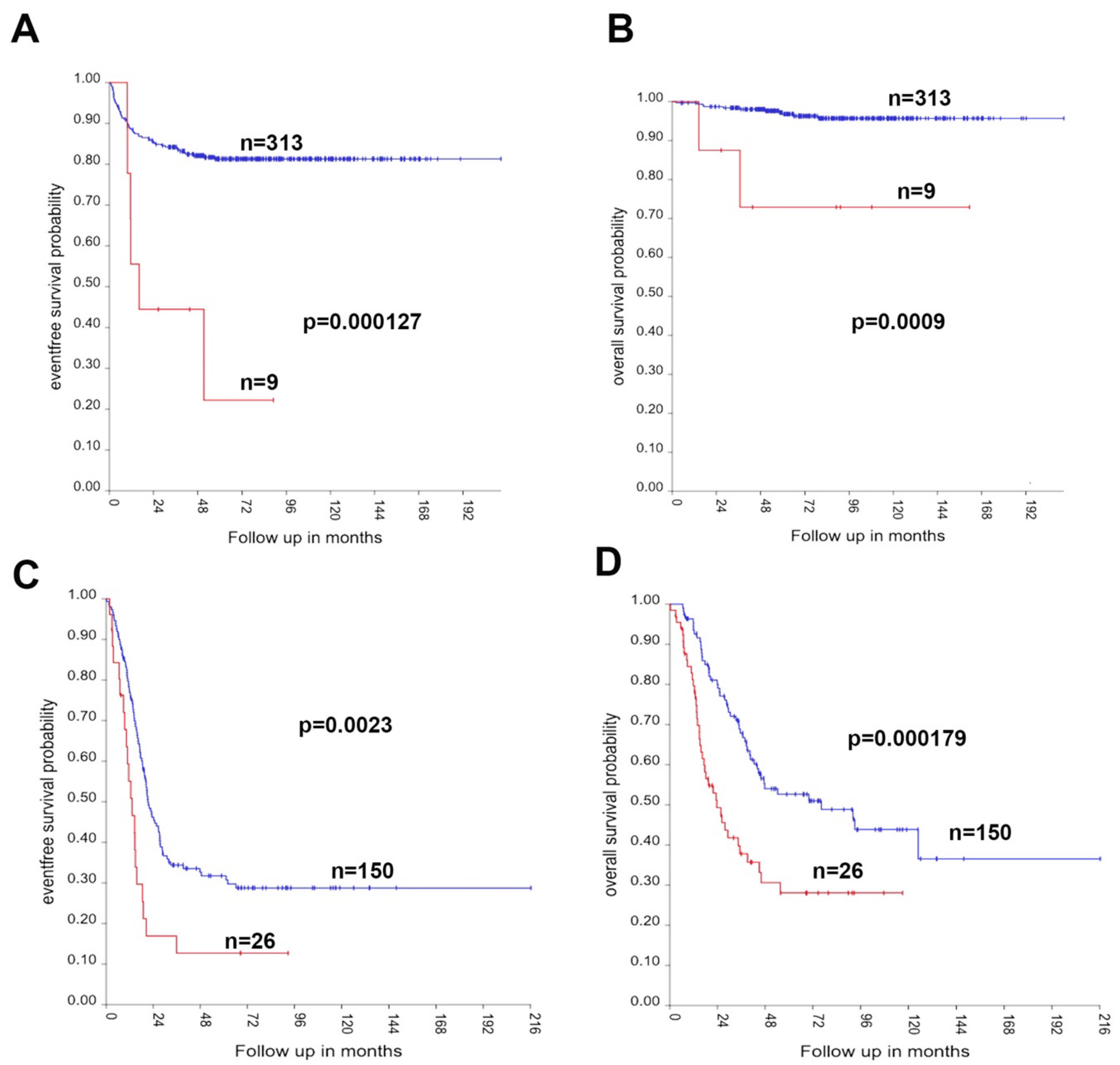


**Figure S3. Association of JMJD6 with survival in low-risk and high-risk patients. (A, B)** Kaplan-Meier survival curve showing that the expression of JMJD6 gene is correlated with worse event-free survival (A) and overall survival (B) in the low-risk patients (SEQC dataset). **(C, D**) Kaplan-Meier survival curve showing that the expression of 17q essential gene is correlated with worse event-free survival (C) and overall survival (D) in the high-risk patients (SEQC dataset).

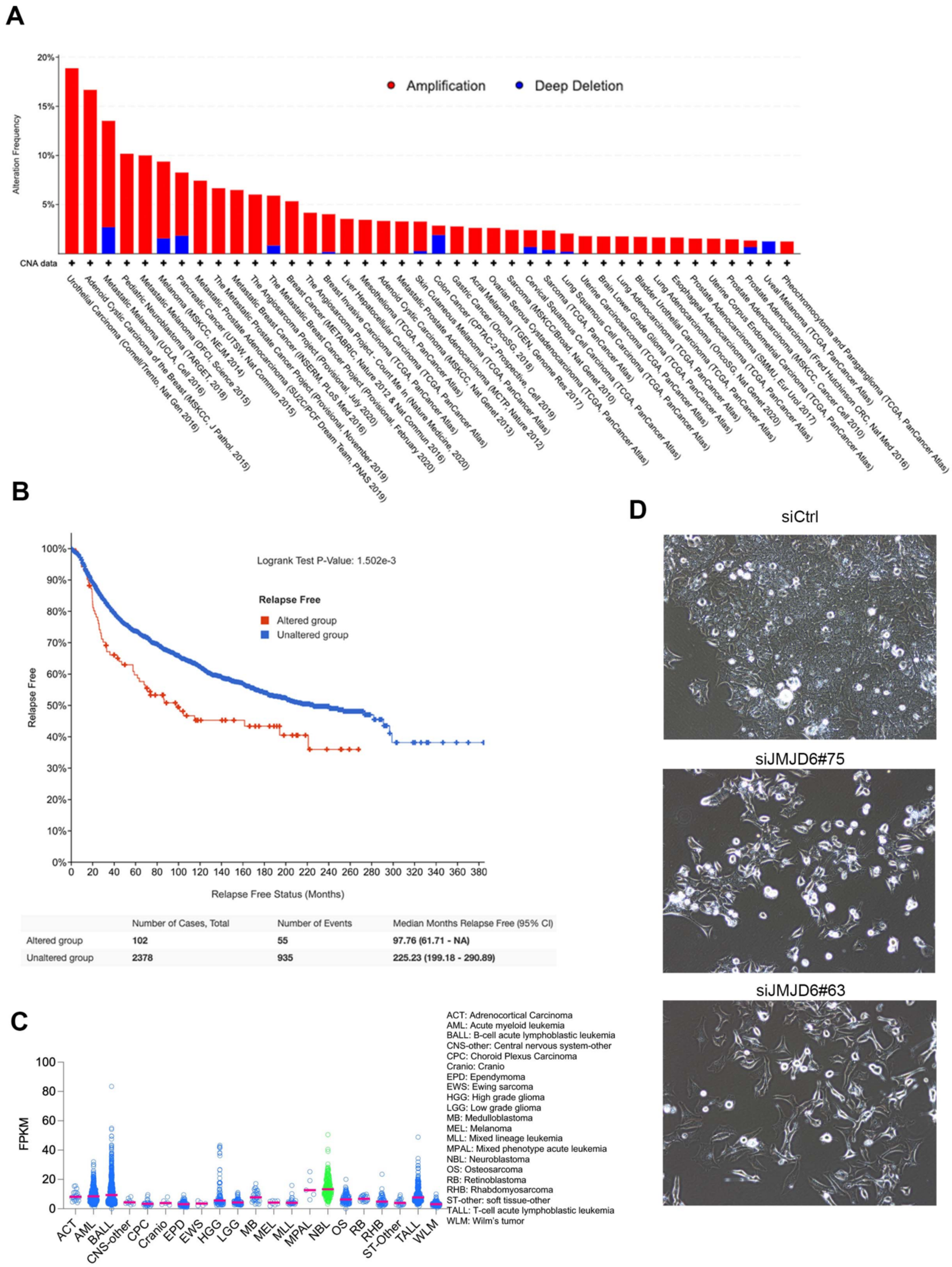


**Figure S4. JMJD6 is amplified in cancers, highly expressed in neuroblastoma.** (**A**) Copy number gain of JMJD6 across a number of cancer types (data from www.cbioportal.com). (**B**) JMJD6 copy number alteration is associated with worse relapse free survival. (**C**) mRNA expression of JMJD6 across a variety of pediatric cancers. (**D**) Cell morphological change of BE2C cells after JMJD6 knockdown on day 5.

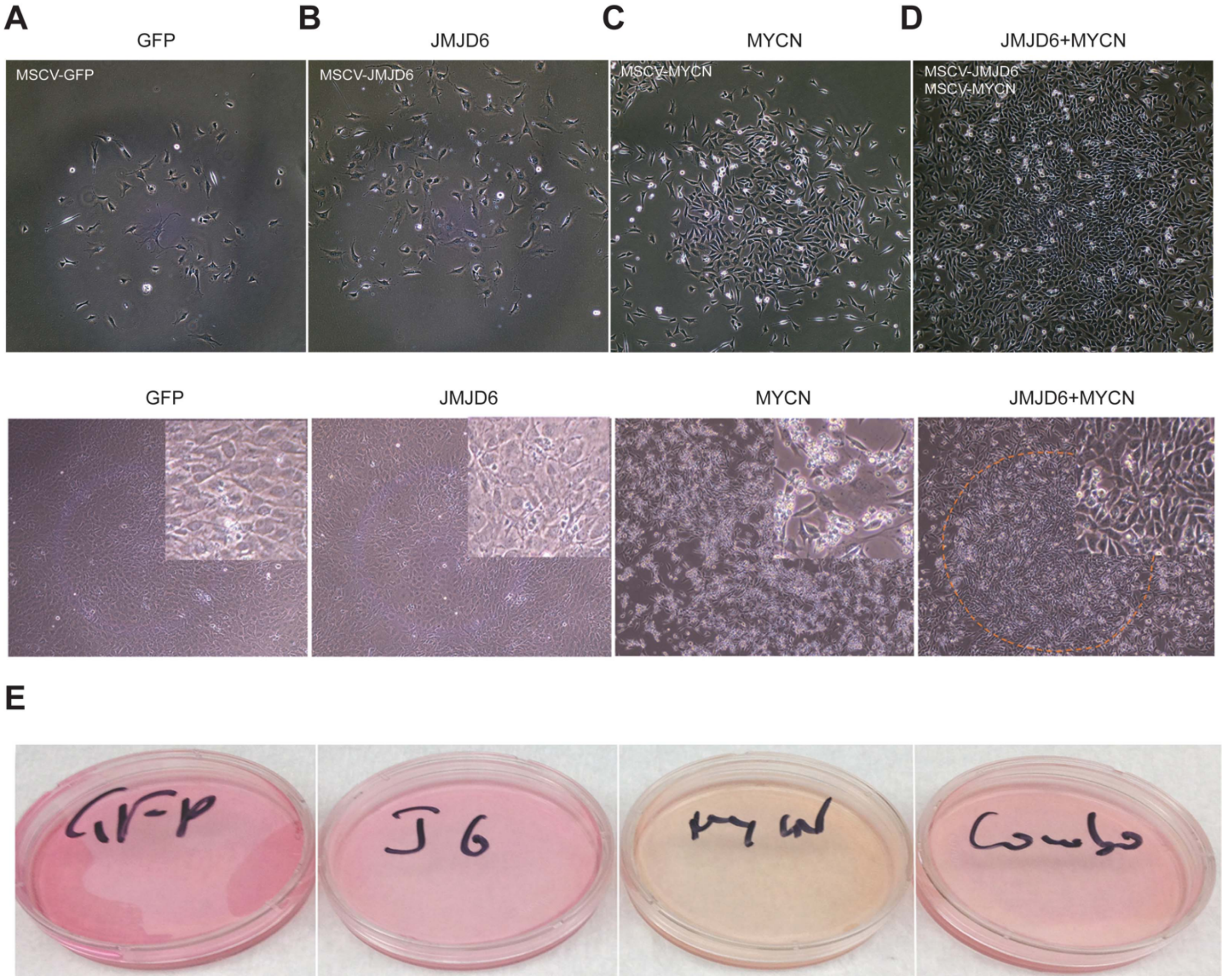


**Figure S5. JMJD6 coordinate with MYC to transform cells.** (**A-D**) The cell morphology after NIH3T3 cells transduced with GFP, JMJD6, MYCN and JMJD6/MYCN before confluence (top) and after confluence (bottom). The inlet showing the higher resolution of images. (**E**) The medium color of NIH3T3 cells transduced with GFP, JMJF6, MYCN and JMJD6/MYCN after they reach confluence.

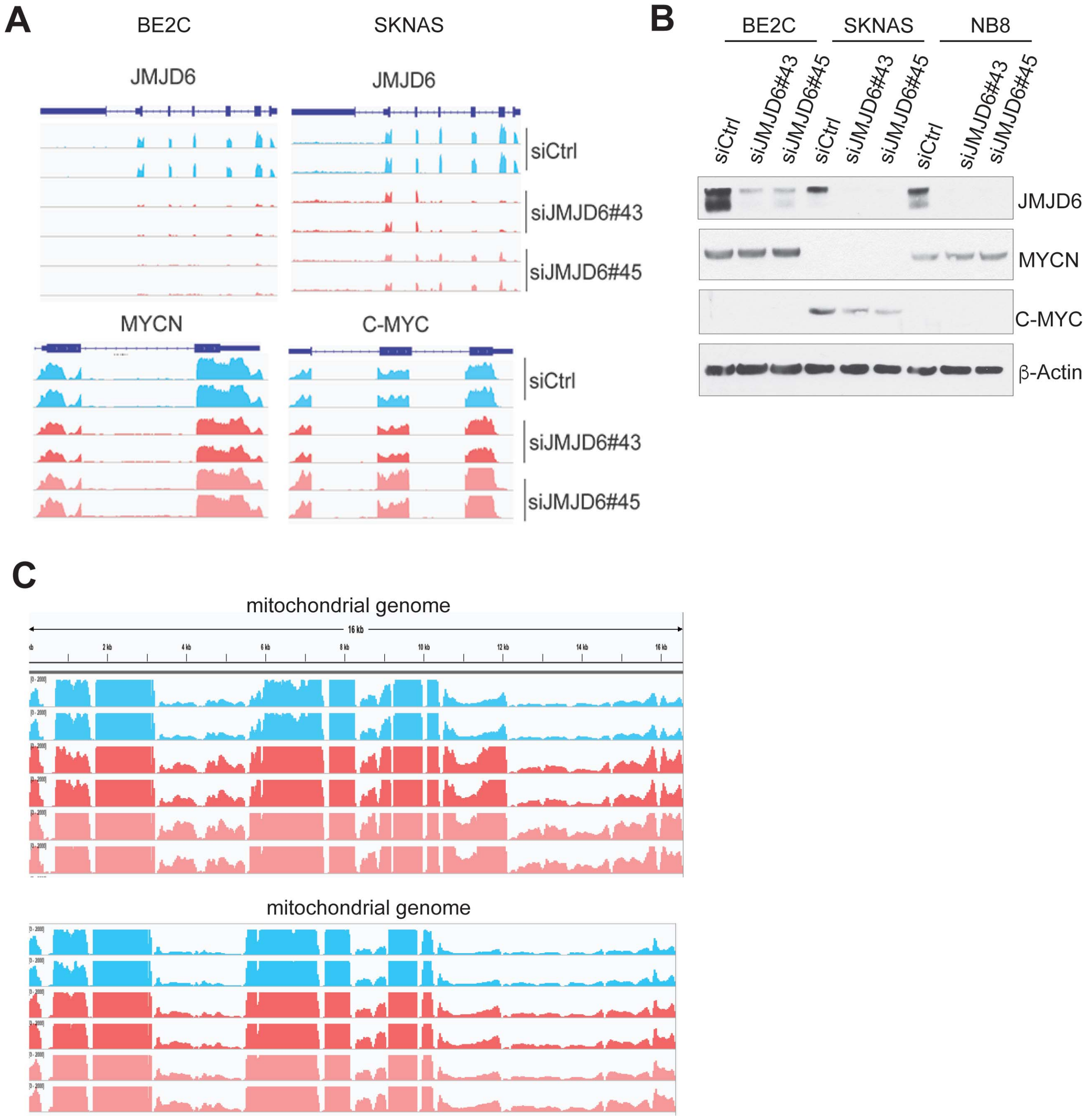


**Figure S6. JMJD6 depletion has no effect on MYC gene transcription but enhances transcription of mitochondrial genome transcription.** (**A**) RNA-seq reads of *JMJD6* and *MYC* (*MYCN/C-MYC*) after JMJD6 knockdown in BE2C and SK-N-AS cells. (**B**) Western blot showing the expression of MYCN and C-MYC after JMJD6 knockdown in BE2C, SK-N-AS and NB8 cells. (**C**) RNA-seq reads of mitochondrial genome genes after JMJD6 knockdown in BE2C and SK-N-AS cells.

**A**

| Gene sets enrichment | SKNAS | | BE2 (C) | |
|---|---|---|---|---|
| | NES | NOM p-val | NES | NOM p-val |
| YU_MYC_TARGETS_UP | -2.57 | 0 | -2.43 | 0.000 |
| ODONNELL_TARGETS_OF_MYC_AND_TFRC_DN | -2.39 | 0 | -2.11 | 0.000 |
| DANG_MYC_TARGETS_UP | -2.12 | 0 | -1.43 | 0.010 |
| CAIRO_PML_TARGETS_BOUND_BY_MYC_UP | -1.81 | 0 | -1.86 | 0.006 |
| MORI_EMU_MYC_LYMPHOMA_BY_ONSET_TIME_UP | -1.73 | 0 | -1.76 | 0.000 |
| DANG_REGULATED_BY_MYC_UP | -1.54 | 0 | N/A | N/A |
| BENPORATH_MYC_TARGETS_WITH_EBOX | -1.49 | 0 | -1.15 | 0.068 |
| FERNANDEZ_BOUND_BY_MYC | -1.45 | 0 | N/A | N/A |
| PID_MYC_ACTIV_PATHWAY | -1.48 | 0.015 | N/A | N/A |
| ALFANO_MYC_TARGETS | -1.17 | 0.036 | N/A | N/A |
| COLLER_MYC_TARGETS_UP | -1.48 | 0.047 | N/A | N/A |

**B**

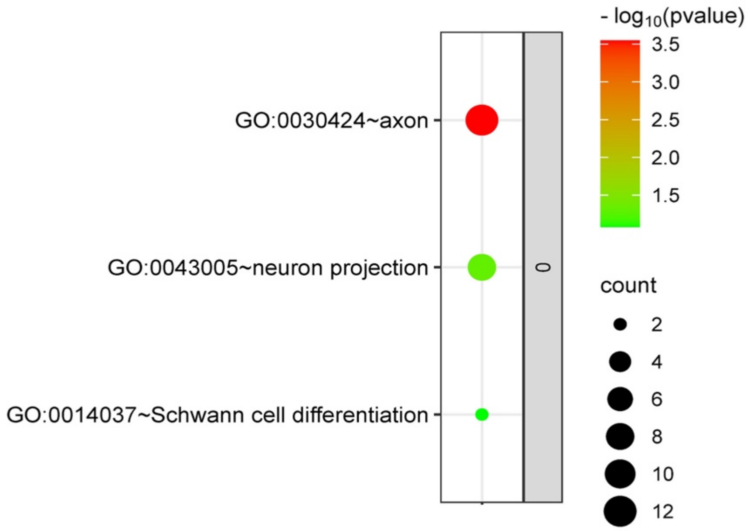


**Figure S7. JMJD6 knockdown downregulates MYC-related pathways and upregulates neuronal differentiation pathways. (A)** Gene set enrichment analysis of RNA-seq data from SK-N-AS and BE2C cells after JMJD6 knockdown. NES = Normalized enrichment score. NOM P value=Nominal P value. **(B)** Bubble plot showing the pathways upregulated in BE2C cells after JMJD6 knockdown.

**A**

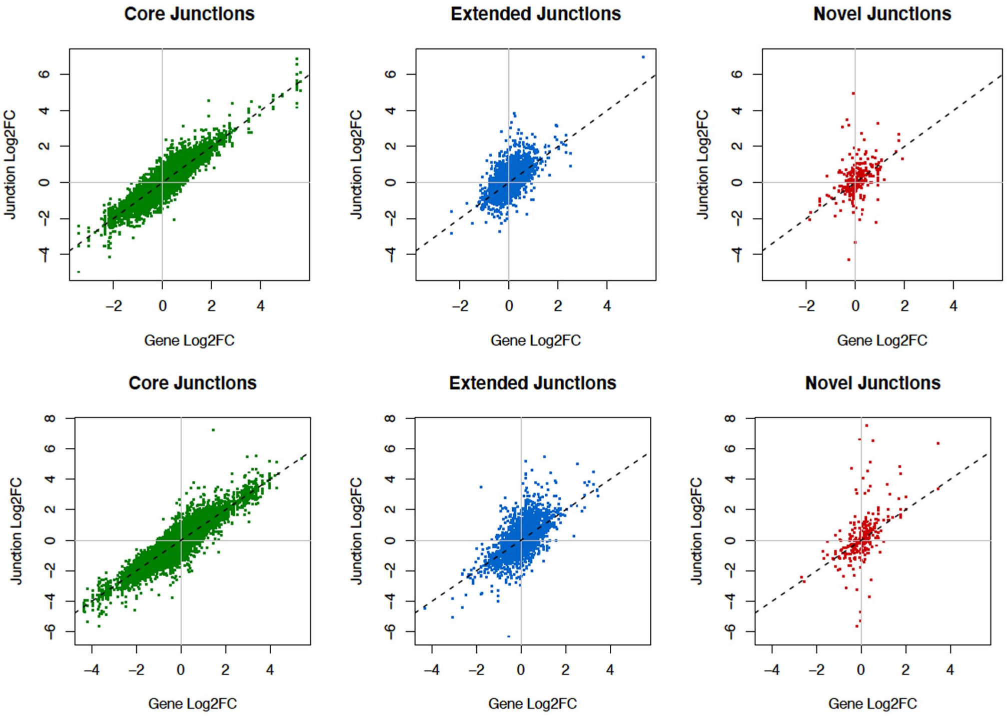


**B**

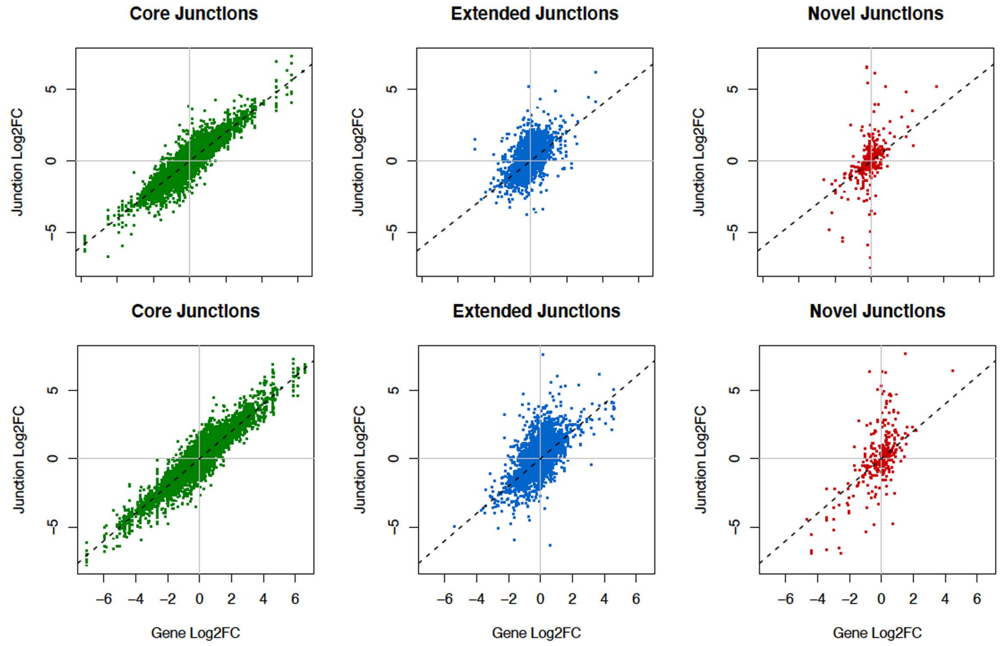


**Figure S8. JMJD6 regulates alternative splicing.** The differentially expressed exon junctions in BE2C (**A**) and (**B**) SK-N-AS cells. Changes in the level of exon junctions compared with the change in gene expression induced by JMJD6 knockdown. Core—defined isoforms identified within the Refseq (NCBI) database; Extended—isoforms defined by the ENSEMBL, AceView, and UCSC databases; Novel—all novel splice junctions that are not detected in untreated samples and are absent from the reference database.

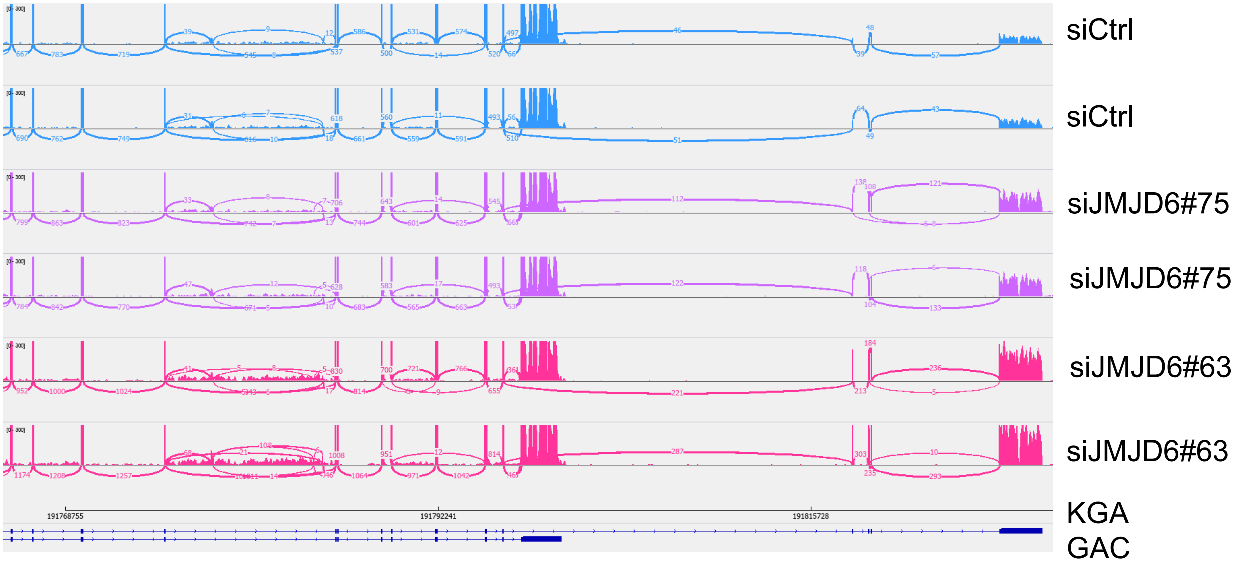


**Figure S9**. **JMJD6 regulates alternative splicing of GLS**. Sashimi plot showing the alternative splicing of *GLS* after JMJD6 knockdown in SKNAS cells in duplicates. The number indicates the RNA-seq read counts of exon junction.

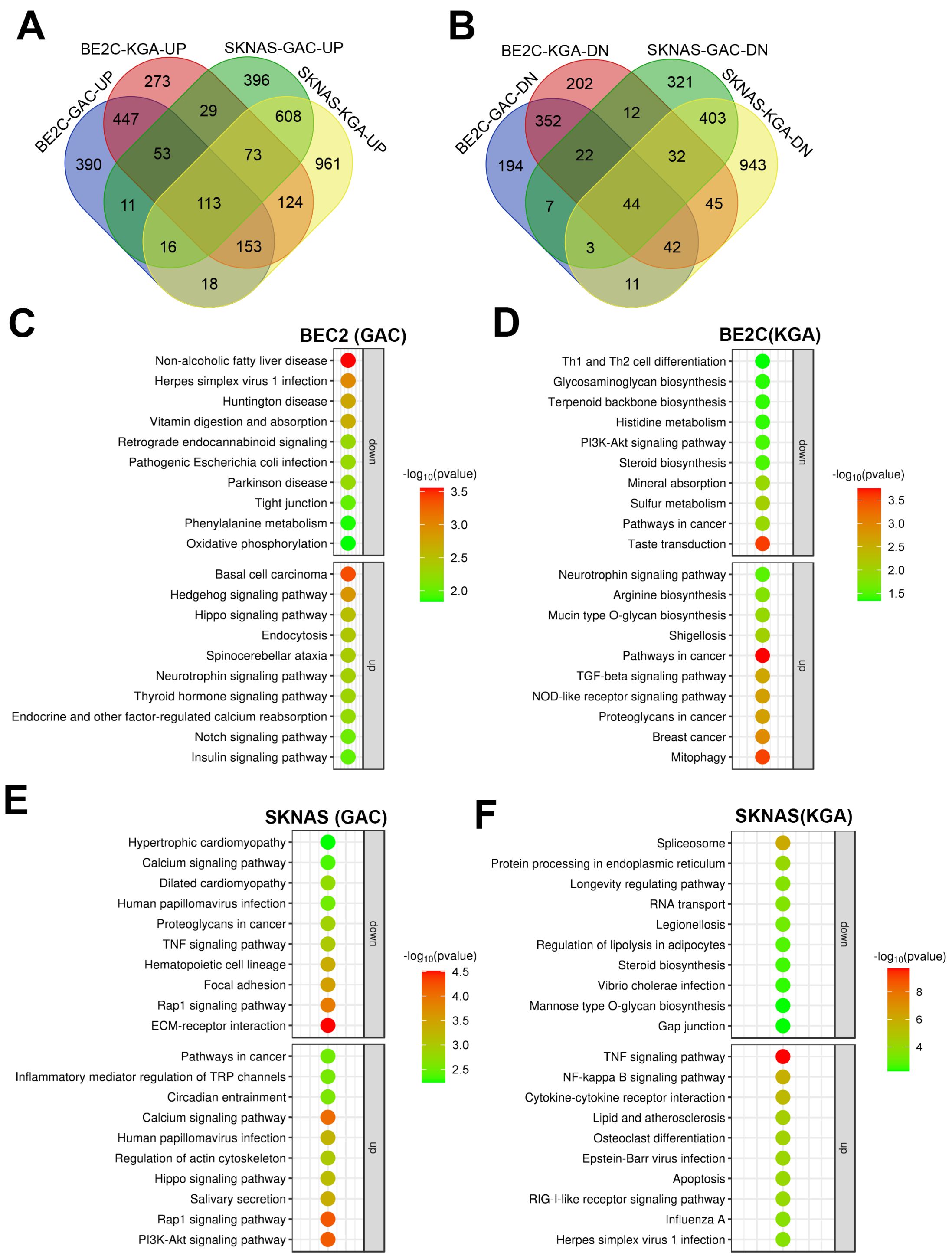


**Figure S10. The differential functions of GAC and KGA in neuroblastoma cells. (A)** Venn diagram showing the genes commonly and differentially upregulated by GAC and KGA in BE2C and SKNAS cells. **(B)** Venn diagram showing the genes commonly and differentially downregulated by GAC and KGA in BE2C and SKNAS cells. **(C-F)** Bubble plots showing the pathways downregulated and upregulated by GAC and KGA in BE2C and SKNAS cells.

**A**

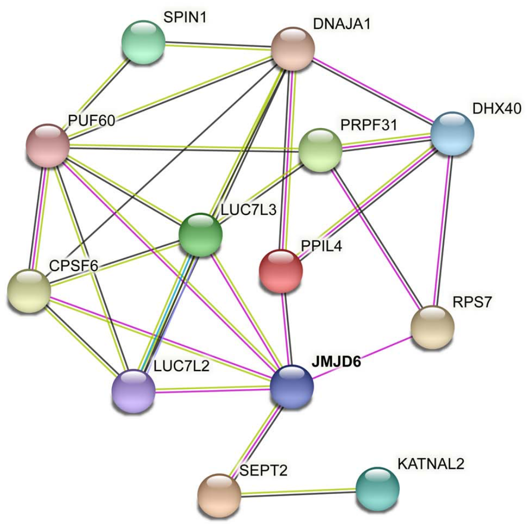


**B**

| Gene Set Name | p-value | FDR q-value |
|---|---|---|
| REACTOME_MRNA_SPLICING | 3.50E-05 | 3.96E-02 |
| REACTOME_METABOLISM_OF_RNA | 6.71E-05 | 3.96E-02 |
| REACTOME_PROCESSING_OF_CAPPED_INTRON_CONTAINING_PRE_MRNA | 7.41E-05 | 3.96E-02 |

**Figure S11. JMJD6 physically interacts with splicing factors.** (**A**) STRING network analysis of JMJD6 interactome in BE2C cells. (**B**) Gene set enrichment analysis of JMJD6 interactome pathway.

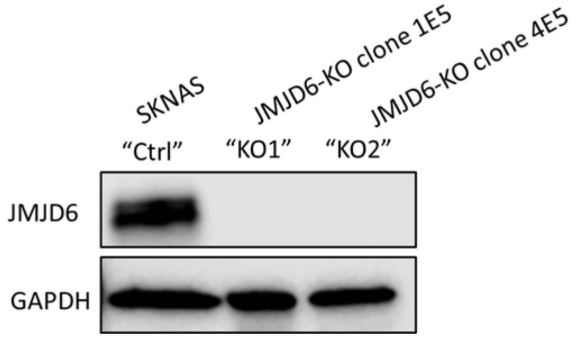


**Figure S12. JMJD6 knockout in SK-N-AS cells.** Western blot showing the complete knockout of JMJD6 in two individual clones.

Table S1. 17q gene list

Table S2. Essential genes in 17q

Table S3. Differentially expressed genes by JMJD6 knockdown

Table S4. Correlation of JMJD6 KO and its co-dependence genes

Table S5. Differential alternative splicing by JMJD6 knockdown

Table S6. GSEA pathways of alternative splicing genes induced by JMJD6 knockdown

Table S7. Mass spectrometry identification of JMJD6 interactome

Table S8. Correlation of metabolite abundance and JMJD6 knockout effect
